# Reverse-engineering human olfactory perception from chemical features of odor molecules

**DOI:** 10.1101/082495

**Authors:** Andreas Keller, Richard C. Gerkin, Yuanfang Guan, Amit Dhurandhar, Gabor Turu, Bence Szalai, Joel D. Mainland, Yusuke Ihara, Chung Wen Yu, Russ Wolfinger, Celine Vens, Leander Schietgat, Kurt De Grave, Raquel Norel, DREAM Olfaction Prediction Challenge Consortium, Gustavo Stolovitzky, Guillermo Cecchi, Leslie B. Vosshall, Pablo Meyer

## Abstract

Despite 25 years of progress in understanding the molecular mechanisms of olfaction, it is still not possible to predict whether a given molecule will have a perceived odor, or what olfactory percept it will produce. To address this stimulus-percept problem for olfaction, we organized the crowd-sourced DREAM Olfaction Prediction Challenge. Working from a large olfactory psychophysical dataset, teams developed machine learning algorithms to predict sensory attributes of molecules based on their chemoinformatic features. The resulting models predicted odor intensity and pleasantness with high accuracy, and also successfully predicted eight semantic descriptors (“garlic”, “fish”, “sweet”, “fruit”, “burnt”, “spices”, “flower”, “sour”). Regularized linear models performed nearly as well as random-forest-based approaches, with a predictive accuracy that closely approaches a key theoretical limit. The models presented here make it possible to predict the perceptual qualities of virtually any molecule with an impressive degree of accuracy to reverse-engineer the smell of a molecule.

**One Sentence Summary:** Results of a crowdsourcing competition show that it is possible to accurately predict and reverse-engineer the smell of a molecule.

## Main Text

In vision and hearing, the wavelength of light and frequency of sound are highly predictive of color and tone. In contrast, it is not currently possible to predict the smell of a molecule from its chemical structure (*1*, *2*). This stimulus-percept problem has been difficult to solve in olfaction because odor stimuli do not vary continuously in stimulus space, and the size and dimensionality of olfactory perceptual space is unknown (*1*, *3*, *4*). Some molecules with very similar chemical structures can be discriminated by humans (*5*, *6*), and molecules with very different structures sometimes produce nearly identical percepts (*2*). Recent computational efforts developed models to relate chemical structure to odor percept (*2*, *7*-*11*), but many relied on psychophysical data from a single 30-year-old study that used odorants with limited structural and perceptual diversity (*12*, *13*).

Twenty-two teams competing in the DREAM Olfaction Prediction Challenge (*14*) were given a large, unpublished psychophysical dataset collected by Keller and Vosshall from 49 individuals who profiled 476 structurally and perceptually diverse molecules, including those that are unfamiliar, unpleasant, or nearly odorless (*15*) (Fig. 1a). We supplied 4884 physicochemical features of each of the molecules smelled by the subjects, including atom types, functional groups, and topological and geometrical properties that were computed using Dragon chemoinformatic software (version 6) (Fig. 1b).

**Fig. 1.**
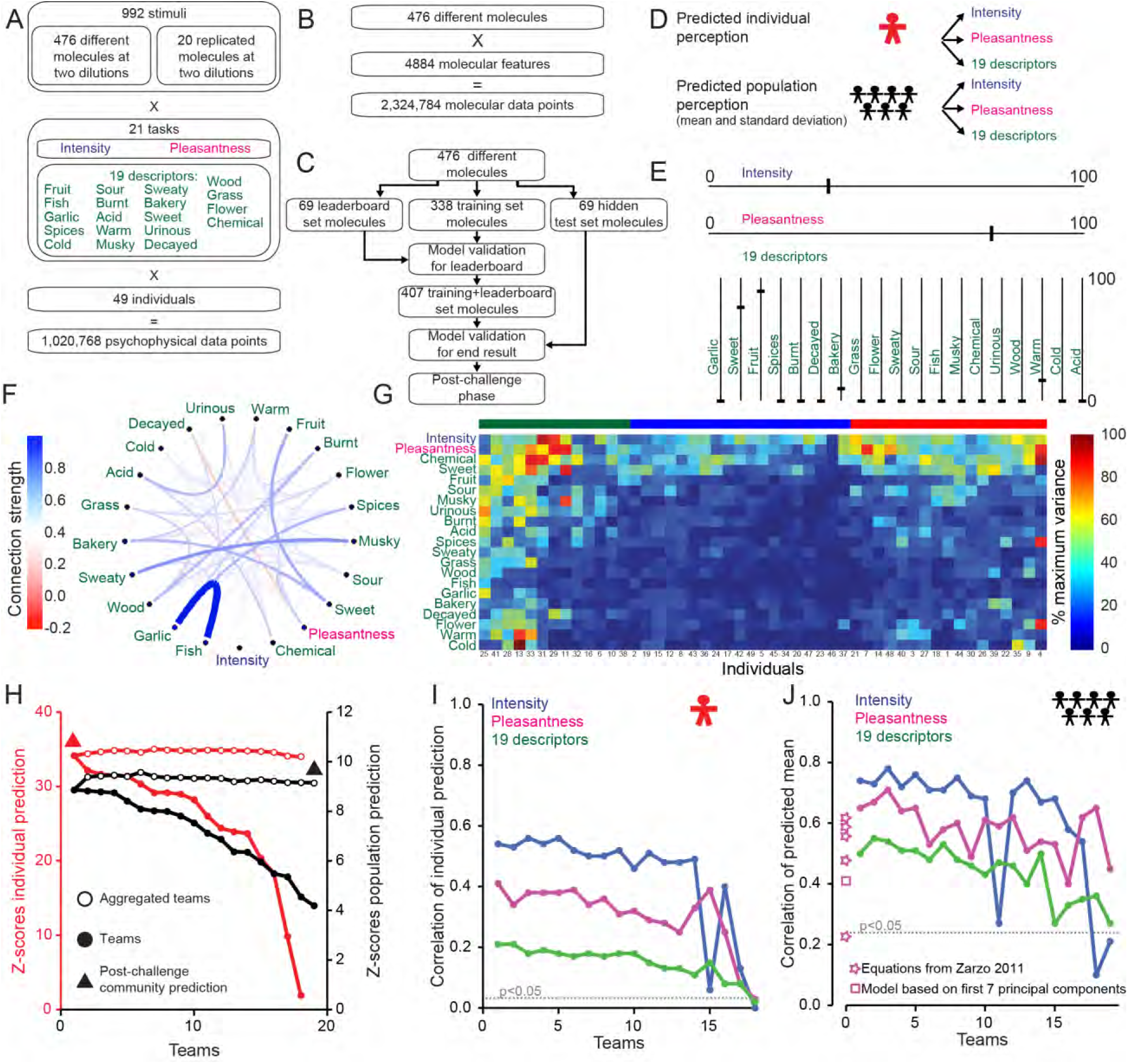
DREAM Olfaction Prediction Challenge. (**A**) Psychophysical data. (**B**) Chemoinformatic data. (**C**) DREAM challenge flowchart. (**D**) Individual and population challenges. (**E**) Hypothetical example of psychophysical profile of a stimulus. (**F**) Connection strength between 21 attributes for all 476 molecules. Width and color of the lines show the normalized strength of the edge. (**G**) Perceptual variance of 21 attributes across 49 individuals for all 476 molecules at both concentrations sorted by Euclidean distance. Three clusters are indicated by green, blue, and red bars above the matrix. (**H**) Model Z-scores, best performers at left. (**I-J**) Correlations of individual (**I**) or population (**J**) perception prediction sorted by team rank. The dotted line represents the p<0.05 significance threshold with respect to random predictions. The performance of four equations for pleasantness prediction suggested by Zarzo (*10*) [from top to bottom: equations (10, 9, 11, 7, 12)] and of a linear model based on the first seven principal components inspired by Khan et al.(*8*) are shown.

Using a baseline linear model developed for the challenge and inspired by previous efforts to model perceptual responses of humans (*8*, *11*), we divided the perceptual data into three sets. Challenge participants were provided with a training set of perceptual data from 338 molecules that they used to build models (Fig. 1c). The organizers used perceptual data from an additional 69 molecules to build a leaderboard to rank performance of participants during the competition. Towards the end of the challenge, the organizers released perceptual data from the 69 leaderboard molecules so that participants could get feedback on their model, and enable refinement with a larger training+leaderboard data set. The remaining 69 molecules were kept as a hidden test set available only to challenge organizers to evaluate the performance of the final models (Fig. 1c). Participants developed models to predict the perceived intensity, pleasantness, and usage of 19 semantic descriptors for each of the 49 individuals and for the mean and standard deviation across the population of these individuals (Fig. 1d-e).

We first examined the structure of the psychophysical data using the inverse of the covariance matrix (*16*) calculated across all molecules as a proxy for connection strength between each of the 21 perceptual attributes (Fig. 1f and Fig. S1). This yielded a number of strong positive interactions including those between “garlic” and “fish”, “musky” and “sweaty”, “sweet” and “bakery” and “fruit”, “acid” and “urinous”, and a negative interaction between pleasantness and “decayed” (Fig. 1f). The perception of intensity had the lowest connectivity to the other 20 attributes. To understand whether a given individual used the full rating scale or a restricted range, we examined subject-level variance across the ratings for all molecules (Fig. 1g). We distinguished three clusters: subjects that responded with high-variance for all 21 attributes (left cluster in green), subjects with high-variance for four attributes (intensity, pleasantness, “chemical”, “sweet”) and either low variance (middle cluster in blue) or intermediate variance (right cluster in red) for the remaining 17 attributes (Figure 1g).

We assessed the performance of models submitted to the DREAM challenge by computing the correlation between the predictions of the 69 hidden test molecules and the actual data. Of the 18 teams who submitted models to predict individual perception, Team GuanLab (author Y.G) was the best performer with a Z-score of 34.18 (Fig. 1h and Data File S1). Team IKW Allstars (author R.C.G.) was the best performer of 19 teams to submit models to predict population perception, with a Z-score of 8.87 (Fig. 1h and Data File S1). The aggregation of all participant models gave Z-scores of 34.02 (individual) and 9.17 (population) (Fig. 1h), and a post-challenge community phase where initial models and additional molecular features were shared across teams gave even better models with Z-scores of 36.45 (individual) and 9.92 (population) (Fig. 1h).

Predictions of intensity were highly correlated with the observed data for both individuals (r=0.56, t-test p<10^-228^) and the population (r=0.78, p<10^-9^) (Fig. 1i, j). Pleasantness was also well predicted for individuals (r=0.41, p<10^-123^) and the population (r=0.71, p<10^-8^) (Fig. 1i, j). The 19 semantic descriptors were more difficult to predict, but the best models performed respectably (individual: r=0.21, p<10^-33^; population: r=0.55, p<10^-5^) (Fig. 1i, j). Previously described models to predict pleasantness (*8*, *10*) performed less well on this dataset than our best model (Fig. 1j). To our knowledge there are no existing models to predict the 19 semantic descriptors.

Random-forest (Fig. 2a and Data File S1) and regularized linear models (Fig. 2b and Data File S1) out-performed other common predictive model types for the prediction of individual and population perception (Fig. 2, Fig. S2, and Data File S1). Although the quality of the best-performing model varied greatly across attributes, it was exceptionally high in some cases (Fig. 2c), and always considerably higher than chance (see dotted line in Fig. 1i), while tracking the observed perceptual values (Fig. S2 for population prediction). In contrast to most previous studies that attempted to predict olfactory perception, these results all reflect predictions of a hidden test set, avoiding the pitfall of inflated correlations due to over-fitting of the experimental data.

**Fig. 2.**
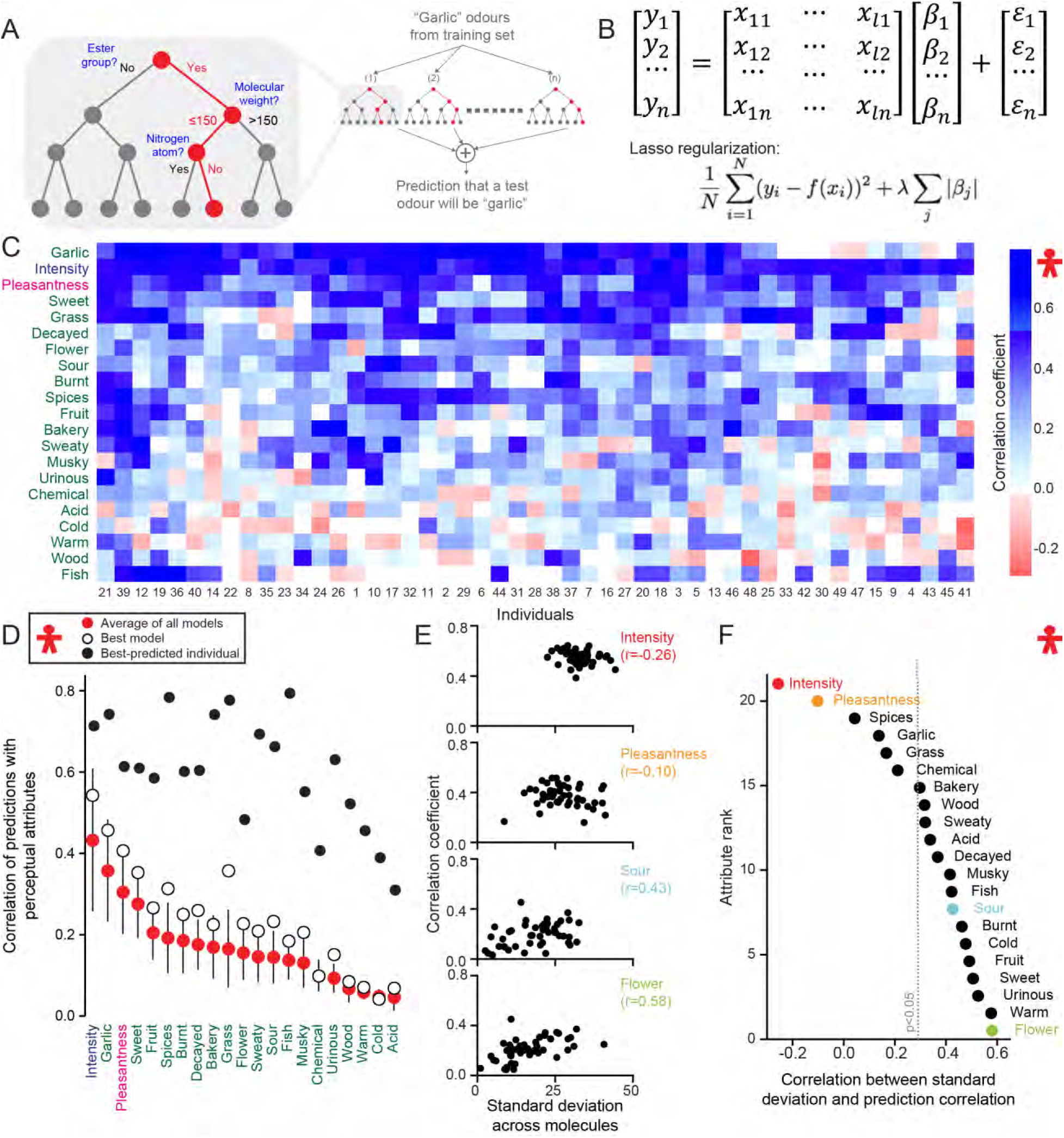
Predictions of individual perception. (**A**) Example of a random-forest algorithm that utilizes a subset of molecules from the training set to match a semantic descriptor (e.g “garlic”) to a subset of molecular features. (**B**) Example of a regularized linear model. For each perceptual attribute y_i_ a linear model utilizes molecular features x_ij_ weighted by β_i_ to predict the psychophysical data of 69 hidden test set molecules, with sparsity enforced by the magnitude of λ. (**C**) Correlation values of best-performer model across 69 hidden test set molecules, sorted by Euclidean distance across 21 perceptual attributes and 49 individuals. (**D**) Correlation values for the average of all models (red dots, mean ± s.d.), best-performing model (white dots), and best-predicted individual (black dots), sorted by the average of all models. (**E**) Prediction correlation of the best-performing random-forest model plotted against measured standard deviation of each subject’s perception across 69 hidden test set molecules for the four indicated attributes. Each dot represents one of 49 individuals. (**F**) Correlation values between prediction correlation and measured standard deviation for 21 perceptual attributes across 49 individuals, color coded as in **E**. The dotted line represents the p<0.05 significance threshold obtained from shuffling individuals.

The accuracy of predictions of individual perception for the best-performing model was highly variable (Fig. 2c), but the correlation of six of the attributes was above 0.3 (Fig. 2d; white circles). The best-predicted individual showed a correlation above 0.5 for 16 of 21 attributes (Fig. 2d). We asked whether the usage of the rating scale (Fig. 1g) could be related to the predictability of each individual. Overall we observed that individuals using a narrow range of attribute ratings, measured across all molecules for a given attribute, were more difficult to predict (Fig. 2e-f, derived from the variance in Figure 1g). The relationship between range and prediction accuracy did not hold for intensity and pleasantness (Fig. 2e-f).

We next compared the results of predicting individual and population perception. The seven best predicted attributes overall (intensity, “garlic”, pleasantness, “sweet”, “fruit”, “spices”, “burnt”) were the same for both individuals and the population (Fig. 2d and Fig. 3a except “fish”). Similarly, the seven attributes that were the most difficult to predict (“acid”, “cold”, “warm”, “wood”, “urinous”, “chemical”, “musky”) were the same for both individual and population predictions (Fig. 2d and Fig. 3a). This suggests some bias in the predictability of more familiar attributes, perhaps due to a better match to a well-defined reference molecule(*15*), and that in this categorization individual perceptions are similar across the population. For the population predictions, the first ten attributes have a correlation above 0.5 (Fig. 3a). The connectivity structure found in Figure 1f follows the model’s performance for the population (Fig. 3a). “Garlic”/“fish”, “sweet”/”fruit” and “musky”/”sweaty” are pairs with high connectivity that were also similarly difficult to predict.

**Fig. 3.**
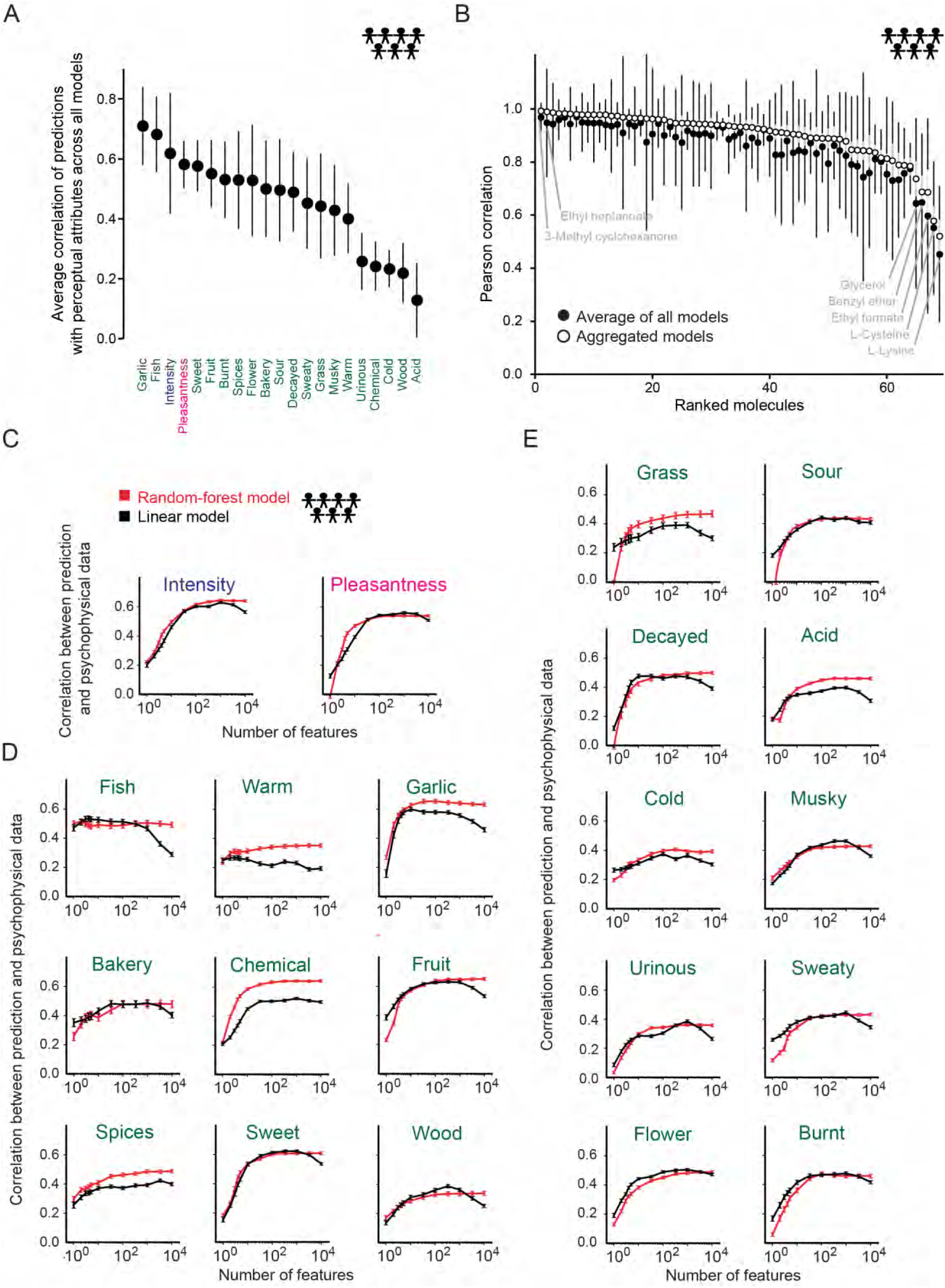
Predictions of population perception. (**A**), Average of correlation of population predictions. Error bars indicate standard deviations calculated across models. (**B**) Ranked prediction correlation for 69 hidden test set molecules produced by aggregated models (open black circles, standard deviation indicated with grey bars) and the average of all models (solid black dots, standard deviation indicated with black bars). (**C-E**) Prediction correlation with increasing number of Dragon features using random-forest (red) or linear (black) models. Attributes are ordered from top to bottom and left to right by the number of features required to obtain 80% of the maximum prediction correlation using the random-forest model. Plotted are intensity and pleasantness (**C**), and attributes that required six or fewer (**D**) or more than six features (**E)**. The combined training+leaderboard set of 407 molecules was randomly partitioned 250 times to obtain error bars for both types of models.

We next analyzed the quality of model predictions for specific molecules in the population (Fig. 3b). The correlation between predicted and observed attributes exceeded 0.9 (t-test p<10^-4^) for 44 of 69 hidden test set molecules when we used aggregated models, and 28 of 69 when we averaged all models (Data File S1). The quality of predictions varied across molecules, but for every molecule the aggregated models exhibited higher correlations (Fig. 3b). The two best-predicted molecules were 3-methyl cyclohexanone followed by ethyl heptanoate. Conversely, the five molecules that were most difficult to predict were L-lysine and L-cysteine, followed by ethyl formate, benzyl ether, and glycerol (Fig. 3b and Fig. S3).

To better understand how the models successfully predicted the different perceptual attributes, we first asked how many molecular features were needed to predict a given population attribute. While some attributes required hundreds of features to be optimally predicted (Fig. 3c-e), both the random-forest and linear models achieved prediction quality of at least 80% of that optimum with far fewer features. By that measure, the algorithm to predict intensity was the most complex, requiring fifteen molecular features to reach the 80% threshold (Fig. 3c). “Fish” was the simplest, requiring only one (Fig. 3d). These results are remarkable because even those attributes needing the most molecular features to predict required only a small fraction of the thousands of chemoinformatic features.

We next asked what features are most important for predicting a given attribute (Fig. 4, Fig. S4, Fig. S5, and Data File S1). The Dragon software calculates a large number of molecular features, but is not exhaustive. In a post-challenge phase (Fig. 1h, triangles), four of the original teams attempted to improve their model predictions by using additional features. These included Morgan (*17*) and NSPDK (*18*), which encode features through the presence or absence of particular substructures in the molecule; experimentally derived partition coefficients from EPI Suite (*19*); and the common names of the molecules. We used cross-validation on the whole dataset to compare the performance of the same models using different subsets of Dragon and these additional molecular features. Only Dragon features combined with Morgan features yielded decisively better results than Dragon features alone both for random-forest (Fig. 5a) and linear (Fig. 5b) models. We then examined how the random-forest model weighted each feature to gain insight into how the models are solving the problem (See Data File S1 for a similar analysis using the linear model). As observed previously, intensity was negatively correlated with molecular size, but was positively correlated with the presence of polar groups, such as phenol, enol, and carboxyl features (Fig. 4a) (*1*, *7*). Predictions of intensity relied primarily on Dragon features.

**Fig. 4.**
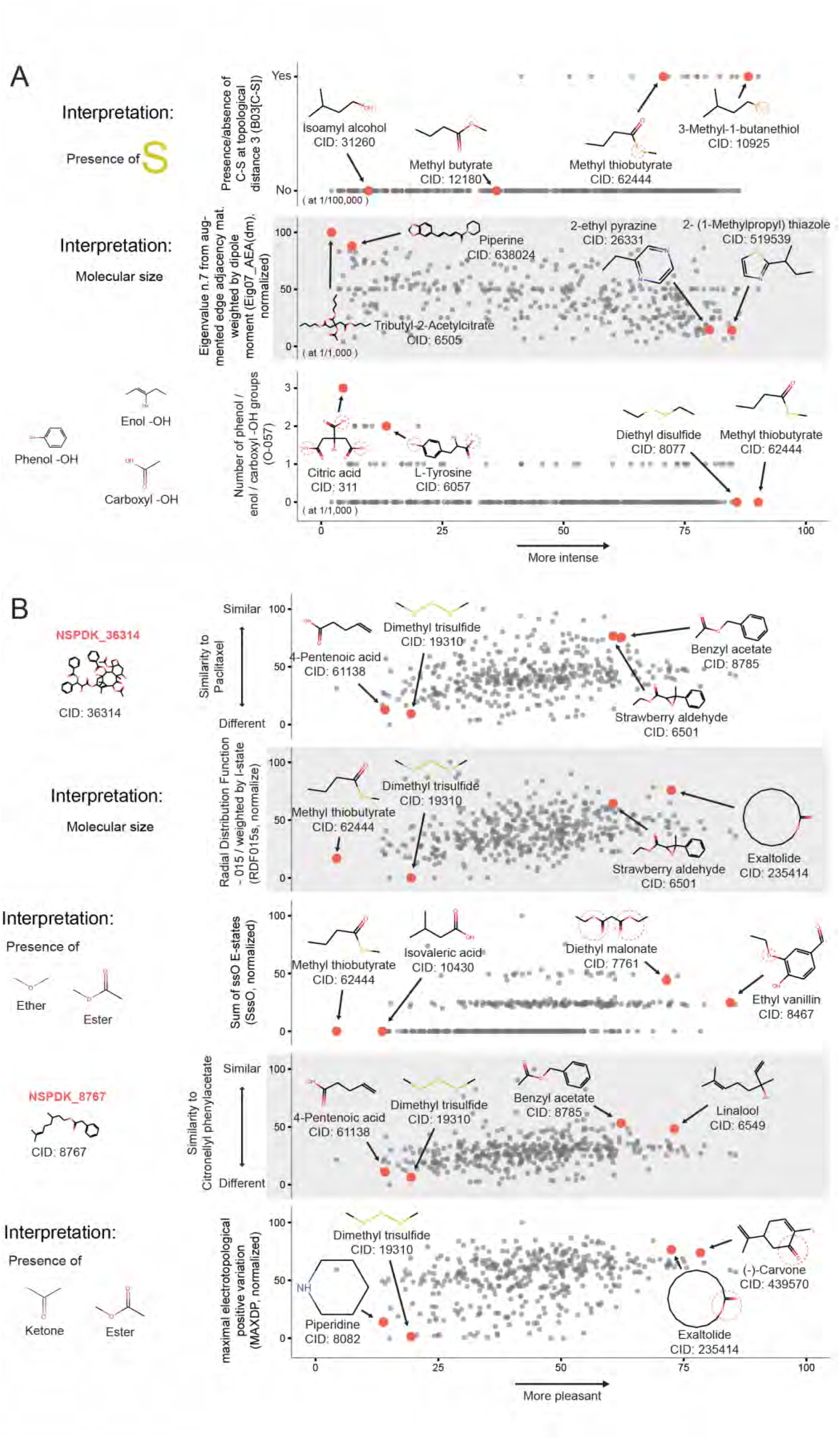
Predicting the smell of specific molecules. (**A-B**), The five most important molecular features selected from Dragon, Morgan, and NSPDK (red text) for predicting (**A**) intensity and (**B**) pleasantness using the random-forest model from the post-challenge phase. Each grey dot represents one of the 407 molecules in the training+leaderboard set, with example molecules indicated by red dots. For (**A**), only three features are shown. The other two are very similar to the one shown in the top panel (B03[C-S]) and the one shown in the middle panel (Eig07_AEA(dm)), respectively.

**Fig. 5.**
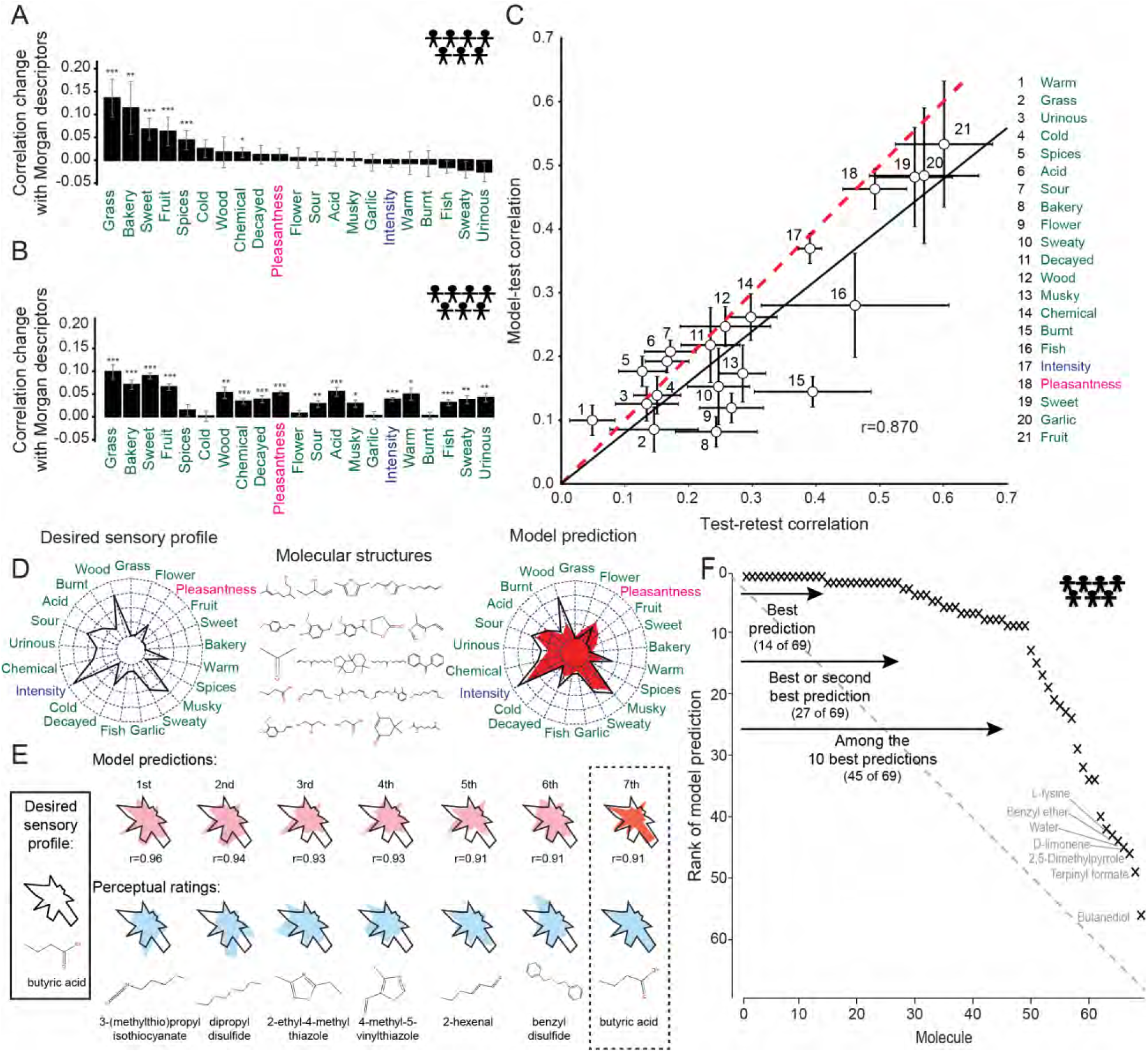
Quality of predictions. (**A-B**) Community phase predictions for random-forest (**A**) and linear (**B**) models using both Morgan and Dragon features for population prediction. The training set was randomly partitioned 250 times to obtain error bars *p<0.05, **p<0.01, ***p<0.001 corrected for multiple comparisons (FDR). (**C)** Comparison between correlation coefficients for model predictions and for test-retest for individual perceptual attributes using the aggregated predictions from linear and random-forest models. Error bars reflect standard error obtained from jackknife resampling of the retested molecules. Linear regression of the model-test correlation coefficients against the test-retest correlation coefficients yields a slope of 0.80 ± 0.02 and a correlation of r=0.870 (black line) compared to a theoretically optimal model (perfect prediction given intra-individual variability, dashed red line). Only the model-test correlation coefficient for “burnt” (15) was statistically distinguishable from the corresponding test-retest coefficient (p<0.05 with FDR correction). (**D**) Schematic for reverse-engineering a desired sensory profile from molecular features. The model was presented with the experimental sensory profile of a molecule (spider plot, left) and tasked with searching through 69 hidden test set molecules (middle) to find the best match (right, model prediction in red). Spider plots represent perceptual data for all 21 attributes, with the lowest rating at the center and highest at the outside of the circle. (**E**) Example where the model selected a molecule with a sensory profile 7^th^ closest to the target, butyric acid. (**F**) Population prediction quality for the 69 molecules in the hidden test set when all 19 models are aggregated. The overall area under the curve (AUC) for the prediction is 0.83, compared to 0.5 for a random model (grey dotted line) and 1.0 for a perfect model.

There is already anecdotal evidence that some chemical features are associated with a sensory attribute. For example, sulfurous molecules are known to smell “garlic” or “burnt”, but no quantitative model exists to confirm this. Our model confirms that the presence of sulfur correlated positively with both “burnt” (Fig. S4a) and "garlic" (Data File S1). Pleasantness was predicted most accurately using a mix of both Dragon and Morgan/NSPDK features. For example, pleasantness correlated with both molecular size (*9*), and similarity to paclitaxel and citronellyl phenylacetate (Fig. 4b). “Bakery” predictions were heavily driven by similarity to the molecule vanillin (Fig. S4b). Morgan features improved prediction in part by enabling a model to template-match target molecules against reference molecules for which the training set contains perceptual data. Thus, structural similarity to vanillin or ethyl vanillin predicts “bakery” without recourse to structural features.

Twenty of the molecules in the training set were rated twice (“test” and “retest”) by each individual, providing an estimate of within-individual variability for the same stimulus. This within-individual variability places an upper limit on the expected accuracy of the optimal predictive model. We calculated the test-retest correlation across individuals and molecules for each perceptual attribute. This value of the observed correlation provides an upper limit to any model, since no model prediction should produce a better correlation than data from an independent trial with an identical stimulus and individual. To examine the performance of our model compared to the theoretically best model, we calculated a correlation coefficient between the prediction of a top-performing random-forest model and the test data. All attributes except “burnt” were statistically indistinguishable from the test-retest correlation coefficients evaluated at the individual-level (Fig. 5c). The slope for the best linear fit of the test-retest and model-test correlation coefficients was 0.80 ± 0.02, with a slope of 1 expected for optimal performance (Fig. 5c). Similar results were obtained using model-retest correlation. Thus, given this dataset, performance of the model is close to that of the theoretically optimal model.

Finally, we evaluated the specificity of the predictions of the aggregated model by calculating how frequently the predicted sensory profile had a better correlation with the actual sensory profile of the target molecule than it did with the sensory profiles of any of the other 68 molecules in the hidden test set (Fig. 5d-e). For 14 of 69 molecules, the highest correlation coincided with the actual sensory profile (p<10^-11^). For an additional 20% it was second highest and 65% of the molecules ranked in the top ten predictions (Fig. 5f and Data File S1; AUC=0.83). The specificity of the aggregated model shows that its predictions could be used to reverse-engineer a desired sensory profile using a combination of molecular features to synthesize a designed molecule.

The DREAM Olfaction Prediction Challenge has yielded models that generated high-quality personalized perceptual predictions. This work significantly expands on previous modelling efforts (*2*, *3*, *7*-*11*) because it predicts not only pleasantness and intensity, but also 19 semantic descriptors of odor quality. The predictive models closely approach the theoretical limits of accuracy when accounting for within-individual variability, and enable the reverse-engineering of a desired perceptual profile to identify suitable molecules from vast databases of chemical structures. Although the current models can only be used to predict the 21 attributes, the same approach could be applied to a psychophysical dataset that measured any desired sensory attribute (e.g. “rose”, “sandalwood”, or “citrus”). How can the highly predictive models presented here be further improved? Recognizing the inherent limits of using semantic descriptors for odors (*13*, *15*, *20*), alternative perceptual data such as ratings of stimulus similarity will be important (*11*). Results of the DREAM Olfaction Prediction Challenge may accelerate efforts to understand basic mechanisms of ligand-receptor interactions, and to test predictive models of olfactory coding in both humans and animal models. Finally, these models have the potential to streamline the production and evaluation of new molecules by the flavor and fragrance industry.

## Acknowledgments

This research was supported in part by grants from the National Institutes of Health (R01DC013339 to JDM, R01MH106674, R01 EB021711 to R.C.G., UL1RR024143 to Rockefeller University), the Russian Science Foundation (#14-24-00155 to M.D.K.), the Slovenian Research Agency (P2-0209 to B.Z.), the Research Fund KU Leuven (C.V.), and IWT-SBO Nemoa (L.S.). A.K. was supported by a Branco Weiss Science in Society Fellowship. L.B.V. is an investigator of the Howard Hughes Medical Institute. P.C.B. has the support of the Ontario Institute for Cancer Research through funding provided by the Government of Ontario and a Terry Fox Research Institute New Investigator Award and a CIHR New Investigator Award. C.V. acknowledges the Research Fund KU Leuven. K.R. is supported by a Grant from the Council of Scientific and Industrial Research.

## Author Contributions

All authors together interpreted the results, approved the design of the figures and the text, which were prepared by A.K., R.C.G., J.D.M., L.B.V., and P.M. LBV is a member of the scientific advisory board of International Flavors & Fragrances, Inc. (IFF) and receives compensation for these activities. IFF was one of the corporate sponsors of the DREAM Olfaction Prediction Challenge. Y.I. is employed by Ajinomoto Co., Inc. J.D.M. is a member of the scientific advisory board of Aromyx and receives compensation for these activities.

## Supplementary Materials

### Materials and Methods

#### Perceptual Data

The psychophysical data for this project were collected between February 2013 and July 2014 as part of the Rockefeller University Smell Study. Data from 49 individuals (28 women, median age 36) were used for the DREAM challenge. The dataset represents a subset of that presented in the original study (*1*), which was unpublished until the DREAM challenge was completed in early 2016. Six individuals declined permission to have their data used in the DREAM challenge. We excluded data on familiarity and edibility ratings for all stimuli, as well as data about whether the individual recognized the smell and how they described it in their own words, as well as data from 4 molecules [compound identification number (CID) 6202: thiamine hydrochloride; CID 24203: sodium phosphate dibasic; CID 2537: camphor; CID 10644: 2-methoxy-3(5 or 6)-isopropylpyrazine]. Twenty-four individuals self-identified as Black, 14 as White, 5 as Asian, and 2 as Native American. Nine individuals self-identified as Hispanic. Individuals provided perceptual ratings of 992 stimuli, 476 different monomolecular chemicals at two different concentrations with 20 molecules tested twice.

Each molecule was presented to individuals at two different concentrations, diluted in paraffin oil so that the "high" and “low” concentrations for each molecule were empirically set to about equal intensity. While molecules were obtained at high purity (>97%), we cannot exclude the possibility that trace contaminants or degradation products account for or add to the odor of the molecule. In the DREAM challenge, teams were asked for predictions of pleasantness and the 19 descriptors only for the “high” concentrations. Individuals were asked to rate each stimulus using 21 perceptual attributes (intensity, pleasantness, and 19 semantic descriptors), by moving an unlabeled slider. The default location of the slider was 50 for intensity and pleasantness, and 0 for the 19 descriptors. For each task, the final position of the slider was translated into a scale from 0 to 100, where 100 signified highest intensity and pleasantness, and the best match of a descriptor for a given stimulus. Further details on the psychophysical procedures and all raw data are available in the original study (*1*).

#### Molecular features

We provided challenge participants with the CID for each molecule, useful for PubChem (https://pubchem.ncbi.nlm.nih.gov/) or other database searches. We used the Dragon software package (version 6; http://www.talete.mi.it) to generate a large number of chemical features for each molecule and made these available to participants.

#### Baseline model for splitting data for the challenge

We developed a linear model with a second layer cubic correction based on a PCA-reduced version of Dragon features to predict the perception of the population. The underlying methodology was used to solve the population prediction and is a multi-linear regression for each of the attributes based on the responses of all individuals and the molecular features of each molecule. The only pre-processing of the data we did was dimensionality reduction of the number of Dragon features, and a log transformation of the values. Based on the above, we chose a random partition that yields good predictive accuracy. We chose the partitions for the leaderboard set and hidden test set based on the distribution of median correlation over test molecules obtained with the model, for different random partitions. The median correlation across molecules selected for the selected partition is above 0.

#### Models

A graphical illustration of one of many decision trees generated by the random-forest algorithm as it evaluates how different structural and physical components determine “garlic” smell is shown in Figure 2a. In each tree, the training data are sequentially partitioned such that each branch point helps increase the accuracy of a prediction. These trees are then aggregated, with their predictions averaged, through a process called bagging. Because the dimensionality of the structural data is high with 4884 Dragon features per molecule and the perception data matrix is sparse, random-forest models are well suited as they help reduce the dimension of the structural data by ignoring unimportant features, and help determine the decision boundary between perceptual ratings of zero and the more informative values. Because most perceptual attributes appeared to depend non-linearly on molecular features, and interactions between features may explain some of the perceptual experience, random-forest models—which can account for these complexities–performed best in this study. However, regularized linear models fared a close second for individual predictions (Data File S1). Linear models (Fig. 2b), which have previously been used to predict perceptual attributes (*2*, *3*), served as a baseline model for the challenge. Their simplicity and good interpretability makes them appealing. Since the number of Dragon features far exceeds the number of molecules, simple linear models such as ordinary least squares regression will produce over-fitting and fail to generalize to untested molecules. Such models will also be sensitive to the highly non-normal distribution of the data and obviously fail to capture non-linear relationships between structural features and perceptual attributes. To overcome these problems, the best linear models used not only the original features, but also their squares (scaled between 0 and 1), and thus were quadratic in the original feature values. To reduce over-fitting, these models used randomized Lasso feature selection, so the summed magnitude of all the regression coefficients is minimized along with the mean-squared error; this automatically selects for models in which many coefficients are zero. Such models were fit on resampled datasets to find the best-fitting and most informative features (Data File S1).

#### Scoring

The training set contained perceptual attribute data from 338 of the 476 molecules. The leaderboard set used for model validation and a hidden test set used for final predictions contained perceptual attribute data from 69 molecules each (Data File S1). Participants had access to the Dragon features for all 476 molecules. However, none of the challenge participants had access to the perceptual attribute data for the 69 molecules in the final hidden test set at any point during the challenge or the community phase. Scoring was handled by the organizers, including PM and RN. Models were scored as follows: for individual prediction, the Pearson correlation between model and data, across test-set molecules, was computed for each individual and attribute. The mean correlations over individuals resulted in 21 attribute-level correlations. These were reduced to (1) the correlation for intensity, (2) the correlation for pleasantness, and (3) the mean of the correlations for the 19 semantic descriptors. These three items were normalized into Z-scores by using the mean and standard deviation for the same dataset with molecule identities shuffled. The final score is the mean of the three Z-scores. Population prediction was scored similarly except that the data were aggregated into means and standard deviations across individuals for each molecule and attribute. Models were asked to predict these means and standard deviations. Here six Z-scores were used, with three corresponding to the means and three to the standard deviations. In both cases we re-scored the models in 1000 bootstraps of the hidden test set.

For individual prediction, the best-performing model remained first in 80 per cent of the bootstrap runs, whereas the second model ranked first in 8 per cent of the runs. For population prediction, the best-performing model remained first in 38 per cent of the bootstrap runs, whereas the second model ranked first in 26 per cent of the runs.

#### Aggregation of models

Participant models were aggregated by first ranking by descending Z-score, then averaging one-by-one following these ranks (the 2 highest ranked models, the 3 highest ranked models, etc.) until all models were aggregated to obtain the same number of aggregations as models.

#### Post-challenge community phase

Five teams (Teams IKW Allstars, GuanLab, KU Leuven, Russ Wolfinger, and Joel Mainland) participated in this phase of the challenge where we discussed ways to enhance the predictions. Each team submitted one new model for both individual and population predictions based on these discussions, which was scored against the same test-set as during the open phase of the challenge. An aggregate model built from these five models was also scored (Figure 1h).

#### Assessing the reverse-engineering of perceptual profiles using the aggregate model

One way to assess the sensitivity of the model’s sensory profile predictions is to calculate the probability of having exactly k correct sensory profile predictions from a list of *n* molecules, that is: 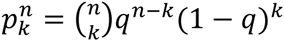 where 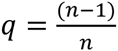 is the probability of matching incorrectly one profile to the list of n molecules.

Here *n*=69 and the aggregated model was able to reverse-engineer k=14 sensory profiles perfectly (20%), so 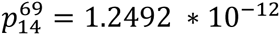.

Another way to measure the performance is to measure the area under the model prediction rank curve (AUC) of Figure 5f. For a perfect model, the prediction rank for every molecule is 1 and so the AUC is the entire plot area: 69*69 (normalized to 1); for a random model all ranks are equally likely and 5f would show a diagonal line (in expectation), with area 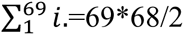 normalized to an AUC of 0.5. For our model presented here has an AUC equal to the perfect model area minus the sum of the ranks of the aggregate model i.e 69*69-830 (normalized to 0.826).

#### Data and Models

Weblinks for data and models are provided below. On the web pages, individual predictions are known as “Subchallenge 1,” and population prediction as “Subchallenge 2.” Model details and code from the best-performing team for individual prediction (Team GuanLab; authors Y.G. and B. P): https://www.synapse.org/#!Synapse:syn3354800/wiki/ (see *files* folder for code)

Model details and code for the best-performing team for population prediction (Team IKW Allstars; author R.C.G.): https://www.synapse.org/#!Synapse:syn3822692/wiki/231036 (check ipython notebooks)

DREAM Olfaction challenge description, participants, leaderboards and datasets: https://www.synapse.org/#!Synapse:syn2811262/wiki/78368

Model descriptions and predictions: https://www.synapse.org/#!Synapse:syn2811262/wiki/78388

Code and details to reproduce analysis for scoring and to reproduce all the analysis for the Figures: http://dream-olfaction.github.io

**Fig. S1.**
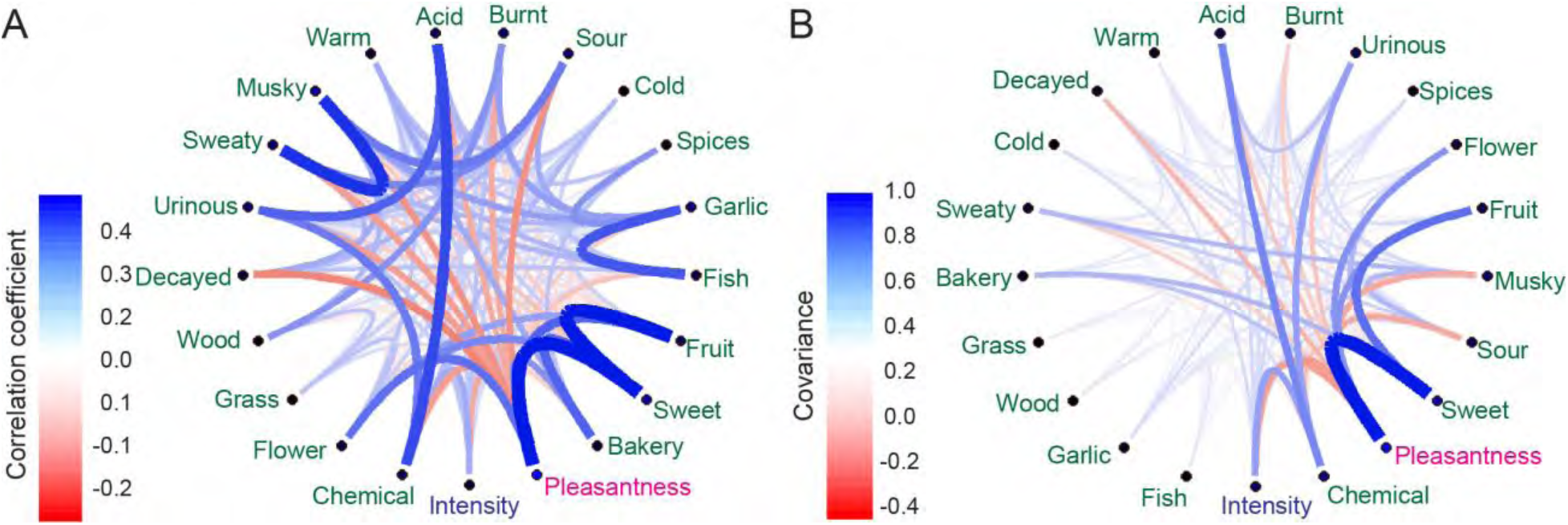
Correlation and covariance of perceptual attributes. (**A-B**) Line width and color represent the strength of the pairwise correlation (**A**) and normalized covariance (**B**) between 21 attributes for all molecules and individuals.

**Fig. S2.**
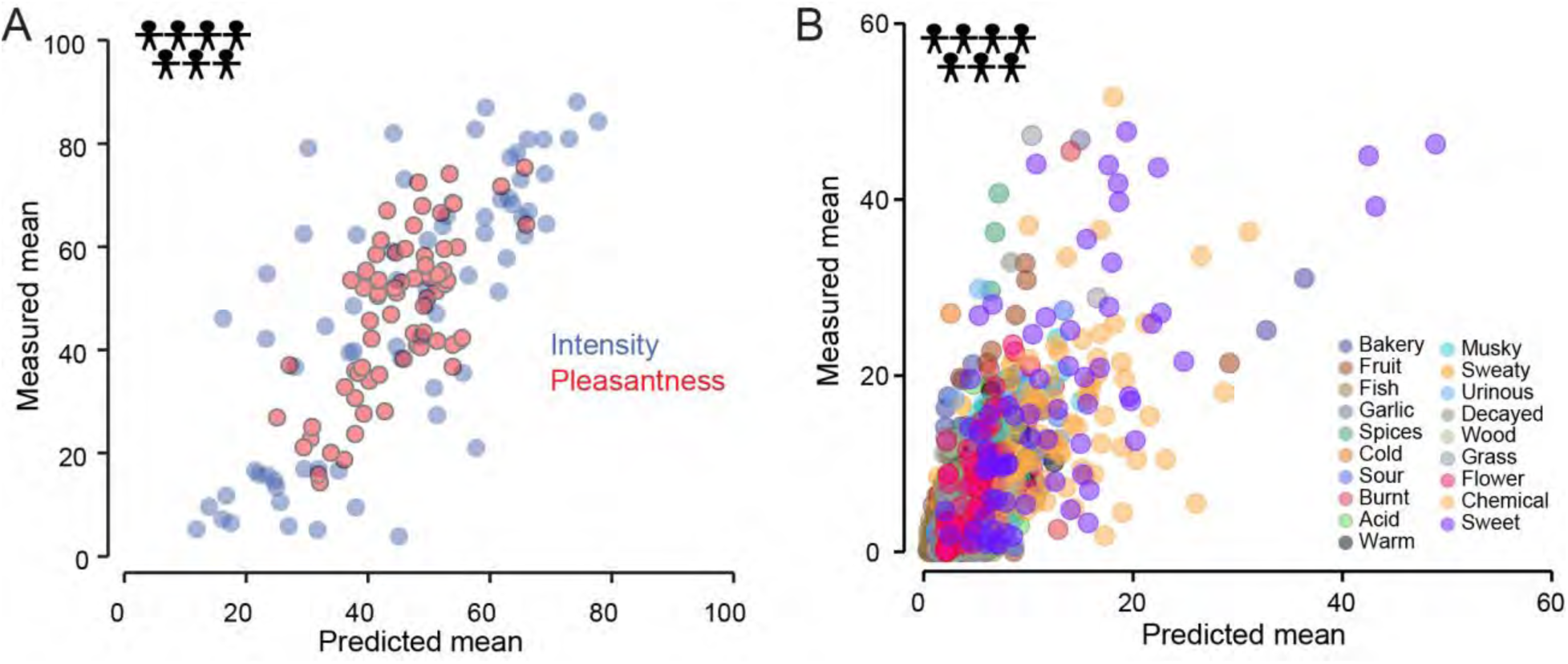
Best performer outcomes for mean and standard deviation for population prediction. (**A-B**) Intensity and pleasantness (**A**) and 19 descriptor (**B**) predictions of the mean of the best-performing team plotted against the observed values for the 69 hidden test set molecules used for model validation.

**Fig. S3.**
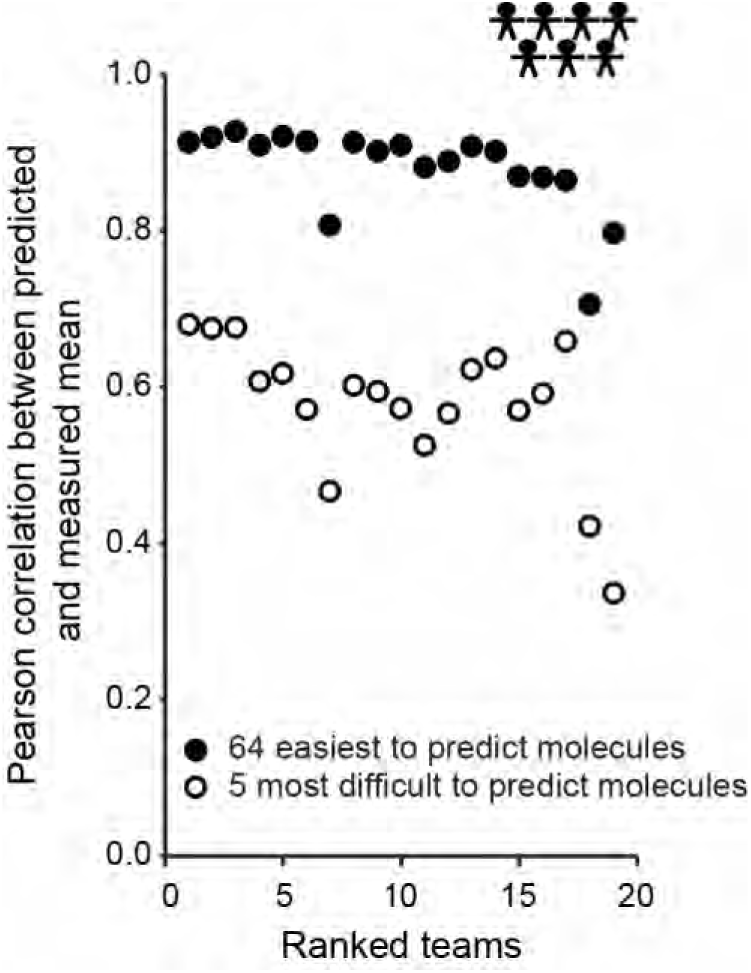
Prediction performance. Pearson correlation between predicted and measured mean perception of the 64 molecules that were the easiest (black dots) and the five molecules that were the most difficult (white dots) to predict. Teams are ordered by their final score for population prediction, with the best performer ranked 1.

**Fig. S4.**
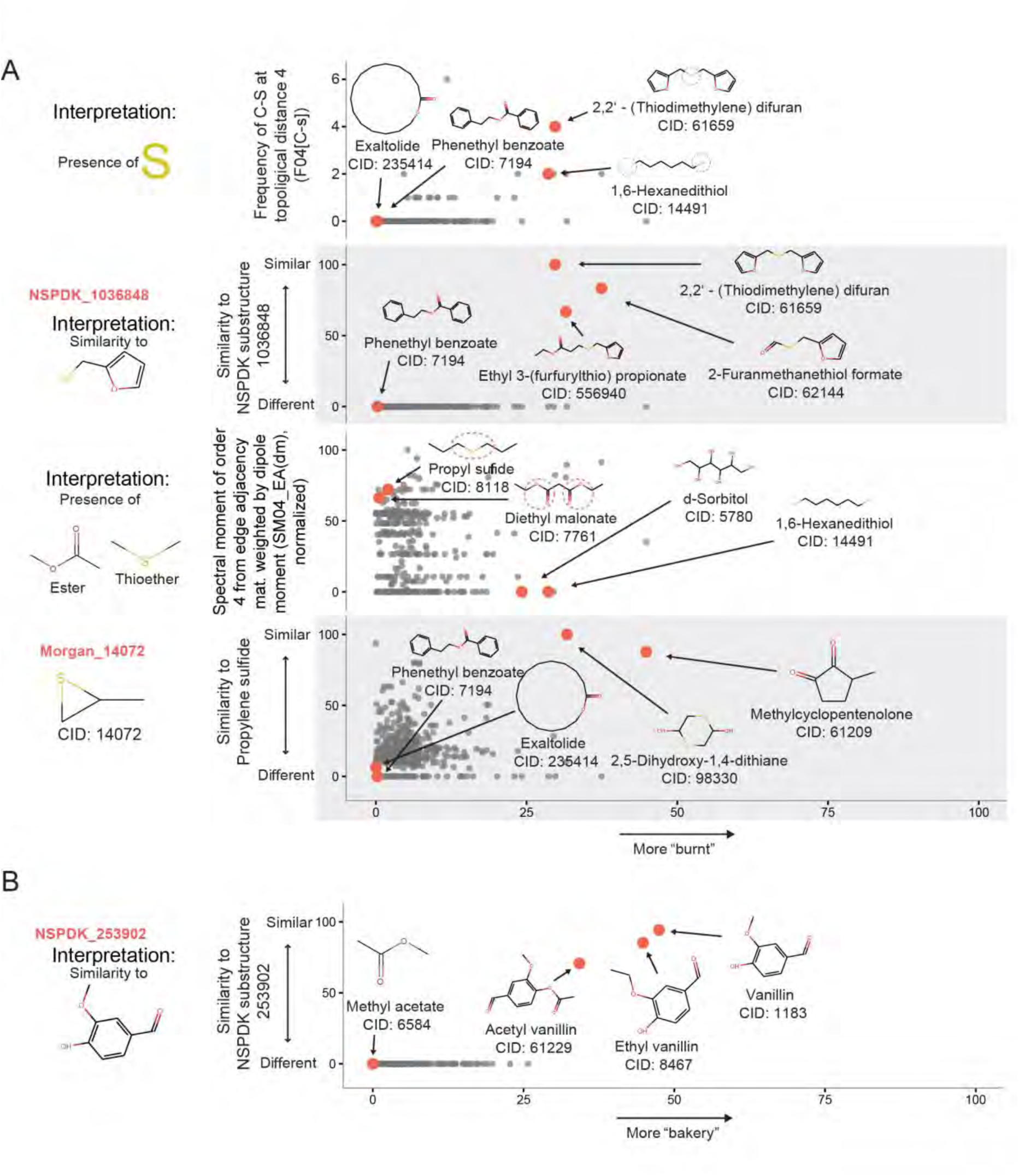
Top molecular features used by the random-forest model from the post-challenge phase as predictors for “burnt” and “bakery.” (**A-B**) Each grey dot represents predictions of each of the 407 molecules in the training+leaderboard set for “burnt” (**A**) and “bakery” (**B**), with example molecules indicated by red dots. In (**A**) only four of the five top features are shown. The fifth feature (R3p+; R maximal autocorrelation of lag 3 / weighted by polarizability) is very similar to the feature depicted in the top panel. In (**B**) only the top feature is shown. The four other features in the top 5 (NSPDK_1022278, NSPDK_722140, NSPDK_250366, NSPDK_555472) are very similar to the one shown.

**Fig. S5.**
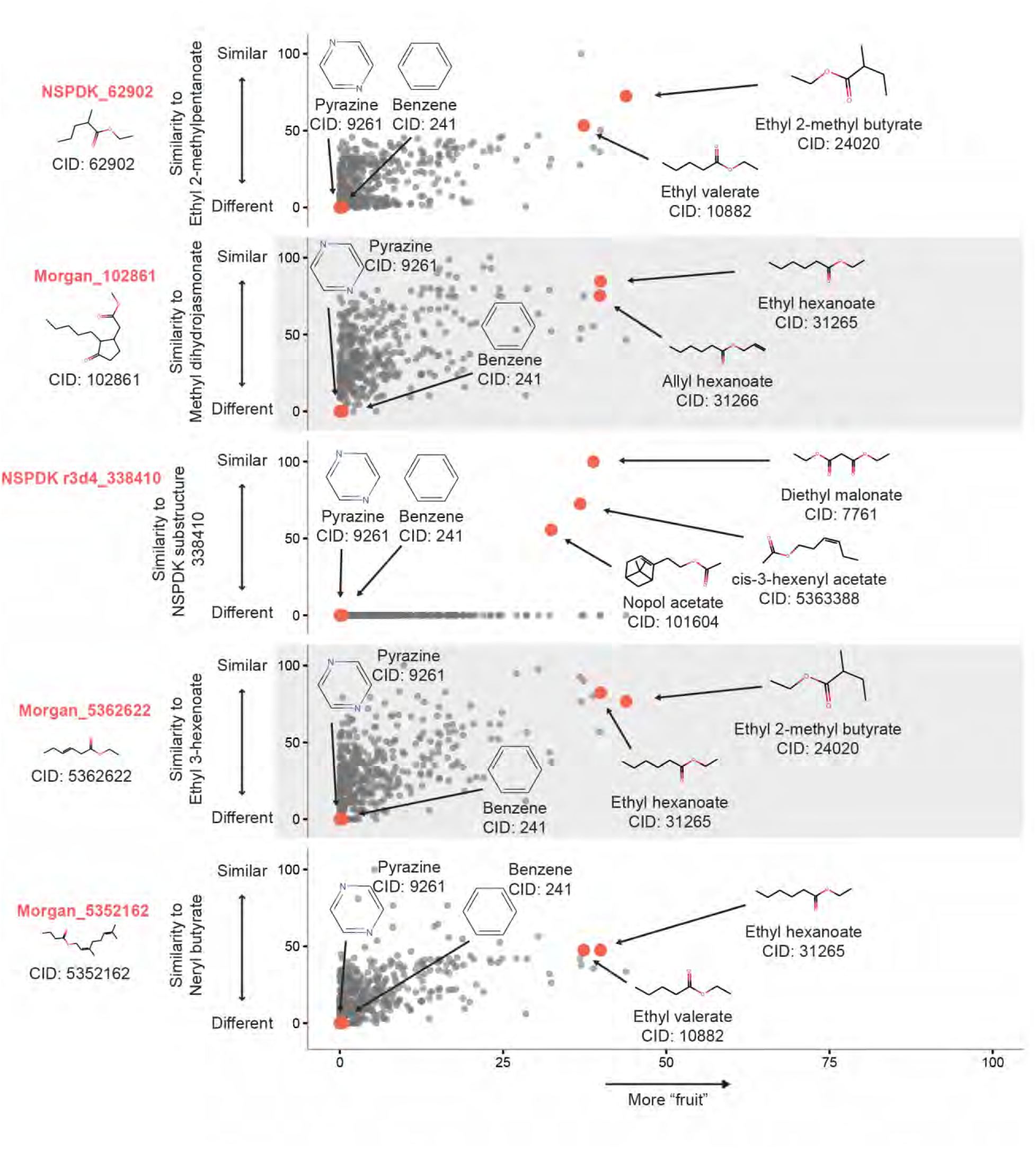
Top 5 molecular features used by the random-forest model from the post-challenge phase as predictors for “fruit”. Each grey dot represents predictions of each of the 407 molecules in the training+leaderboard set for “fruit”, with example molecules indicated by red dots.

**Data File S1. Raw data including prediction scores and methods, correlation values, molecule CIDs and top molecular features.**

